# Loss of β-ketoacyl acyl carrier protein synthase III activity restores multidrug-resistant *Escherichia coli* sensitivity to previously ineffective antibiotics

**DOI:** 10.1101/2022.02.18.481121

**Authors:** Yaoqin Hong, Jilong Qin, Anthony D. Verderosa, Sophia Hawas, Bing Zhang, Mark A. T. Blaskovich, John E. Cronan, Makrina Totsika

## Abstract

Antibiotic resistance is one of the most prominent threats to modern medicine. In the latest World Health Organization list of bacterial pathogens that urgently require new antibiotics, nine out of 12 are Gram-negative, with four being of ‘Critical Priority’. One crucial barrier restricting antibiotic efficacy against Gram-negative bacteria is their unique cell envelope. While fatty acids are a shared constituent of all structural membrane lipids, their biosynthesis pathway in bacteria is distinct from eukaryotes making it an attractive target for new antibiotic development that remains less explored. Here, we interrogated the redundant components of the bacterial Type II <u>F</u>atty <u>A</u>cid <u>S</u>ynthesis (FAS II) pathway, showing that disrupting FAS II homeostasis in *Escherichia coli* through deletion of the *fabH* gene damages the cell envelope of antibiotic susceptible and antibiotic resistant clinical isolates. The *fabH* gene encodes the β-ketoacyl acyl carrier protein synthase III (KAS III), which catalyzes the initial condensation reactions during fatty acid biosynthesis. We show that *fabH* null mutation potentiated the killing of multi-drug resistant *E. coli* by a broad panel of previously ineffective antibiotics, despite the presence of relevant antibiotic resistance determinants, for example, carbapenemase *kpc2*. Enhanced antibiotic sensitivity was additionally demonstrated in the context of eradicating established biofilms and treating established human cell infection *in vitro*. Our findings showcase the potential of FabH as a promising target that could be further explored in the development of therapies that may repurpose currently ineffective antibiotics or rescue failing last-resort antibiotics against Gram-negative pathogens.

**IMPORTANCE:** Gram-negative pathogens are a major concern for global public health due to increasing rates of antibiotic resistance and the lack of new drugs. A major contributing factor towards antibiotic resistance in Gram-negative bacteria is their formidable outer membrane, which acts as a permeability barrier preventing many biologically active antimicrobials from reaching the intracellular targets and thus limiting their efficacy. Fatty acids are the fundamental building blocks of structural membrane lipids and their synthesis constitutes an attractive antimicrobial target as it follows distinct pathways in prokaryotes and eukaryotes. Herein, we identified a component of fatty acid synthesis, FabH, as a ‘gate-keeper’ of outer membrane barrier function. Without FabH, Gram-negative bacteria become susceptible to otherwise impermeable antibiotics and are re-sensitised to killing by last-resort antibiotics. This study supports FabH as a promising target for inhibition in future antimicrobial therapies.

## INTRODUCTION

The outer membrane (OM) of Gram-negative bacteria is critical to their survival within harsh yet fluctuating environments (1). Unlike the canonical cytoplasmic membrane, the OM has an asymmetrical design, in which the inner leaflet is almost exclusively composed of phospholipids (PLs) while the outer leaflet is filled with the lipid A components of lipopolysaccharide (LPS) (2). Each LPS molecule carries anionic charges due to phosphate and carboxylate groups, which provides much of the OM rigidity through intermolecular interactions. The uniformly distributed LPS molecules on the outer leaflet of OM make it a potent barrier and guard the bacterium from harmful compounds (3). Moreover, each LPS molecule is capped by a hydrophilic core oligosaccharide, and in most cases, a terminal long-chain polysaccharide is also attached to the core oligosaccharide. In effect, this creates an additional stablising and protecting water-rich layer extending from the cell surface (4). For nutrient uptake, the otherwise impermeable OM houses a set of specialized porin proteins that allows solute exchange (3).

Over the past decades, the emergence of resistance to currently available antibiotics among many clinically important bacterial pathogens has been recognized as a major threat to global public health. In response to this challenge, the World Health Organization released a list of priority pathogens urgently requiring new antimicrobials (5, 6). Within the Critical Priority Category of this list is the third-generation cephalosporin and/or carbapenem-resistant Enterobacteriaceae (including *Escherichia coli, Klebsiella, Serratia*, and *Proteus*). Many circulating pathogenic *E. coli* lineages are multi-drug resistant (MDR), and therefore, remain susceptible to only a few available treatment options, which are fast diminishing (7, 8). One key contributor to antimicrobial resistance (AMR) in Gram-negative bacteria is their OM. Most effective antibiotics access their intracellular targets using OM-spanning hydrophilic porin channels, but this transport is restricted by both the chemical properties and the size of the permeant antibiotics. In the Enterobacteriaceae family, only small hydrophilic antibiotics (< 600 kDa) can permeate the generalized porins (9, 10).

In bacteria, the essential acyl chains of PL and LPS are *de novo* synthesised by the highly conserved Type II Fatty Acid Synthesis pathway (FAS II) (11, 12). The structural heterogeneity of acyl chains present in membrane lipids can determine how bacteria respond to challenging and fluctuating environments (13-19). As seen in other essential pathways, FAS II has evolved specialized components, which possess overlapping biochemical activities and complex redundancies. Thus, several enzymes are redundant despite the essentiality of their biochemical activities. For example, in *E. coli*, three β-ketoacyl acyl carrier protein (ACP) synthases (KAS proteins), including FabB for KAS I activity (encoded by *fabB*), FabF for KAS II (encoded by *fabF*), and FabH for KAS III (encoded by *fabH*), collectively take parts in the Claisen condensation reactions to drive FA biosynthesis (11). FabH initiates FA biosynthesis through condensing acetyl coenzyme A (CoA) with malonyl ACP (20), while FabB and FabF take overlapping roles in the subsequent polymerisation steps (11). The associated substrate specificity has been reported in great detail by J. L. Garwin et al. (21).

The PL-LPS ratio, maintenance of membrane asymmetry, surface charge profiles of the surface exposed LPS, and chemical differences within both the PLs and LPS components collectively define the overall physicochemical and biological properties of the OM (22). As such, progress in understanding the OM may lead to valuable tactics to circumvent this formidable protective barrier and improve antibiotic cell entry in Gram-negative bacteria. In this work, we examined the role of redundant FAS II genes in maintaining the antimicrobial exclusion properties of the membrane barrier in clinically relevant uropathogenic *E. coli* (UPEC) strains, including a reference MDR isolate from the globally disseminated sequence type 131 (ST131) lineage (23, 24). We report UPEC Δ*fabH* strains lacking KAS III activity display a severely defective membrane envelope and therefore are highly sensitive to antibiotic killing (up to 41-fold reduced MIC), even by otherwise ineffective drugs and while still harboring relevant AMR determinants. Moreover, this increased sensitivity held true in established biofilm eradication and the treatment of infected human bladder cell monolayers. Together, this work showcases FabH as a promising FAS II component that could be therapeutically targeted to rescue failing last-resort antibiotics and expand the range of currently available antibiotic treatments for Gram-negative infections.

## RESULTS

### FabH contributes to outer membrane barrier function in UPEC

To study the involvement of the redundant genes in the FAS II pathway in the maintenance of the OM barrier, we used two model UPEC strains, CFT073 (a drug-sensitive reference pyelonephritis isolate) and EC958 (a reference MDR ST131 cystitis isolate), and constructed null mutants of the *fabH, fabF, fabR, fadR*, and *fadD* genes. Mutants were evaluated for OM defects by measuring their susceptibility to sub-inhibitory concentrations of vancomycin (25). UPEC Δ*fabF*, Δ*fabR*, Δ*fadR*, and Δ*fadD* mutants had similar growth to wild-type (WT) on LB-Lennox containing 50 *µ*g/mL vancomycin, in both strains (Fig 1). Like previous *E. coli* K-12 Δ*fabH* studies (26, 27), UPEC Δ*fabH* strains grew remarkably slower than the WT strains (Supplementary material, Fig S1), and this is reflected in the reduced colony size illustrated in Fig 1A.

**Fig 1.**
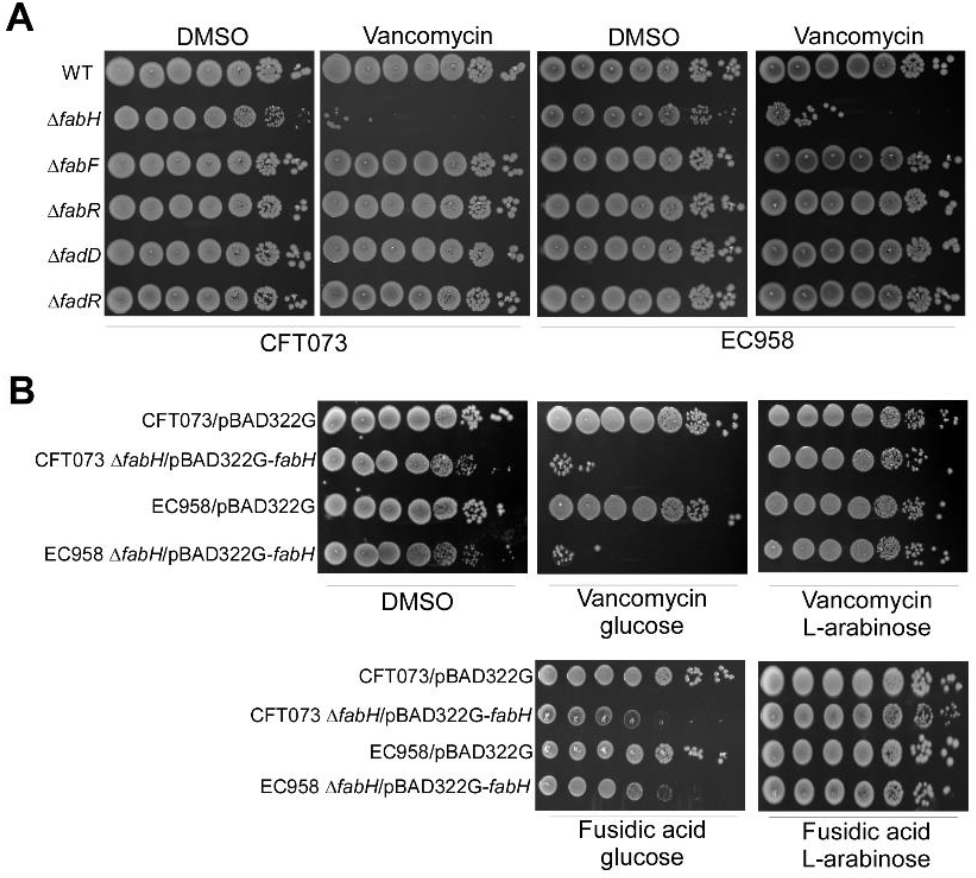
Screening UPEC FAS II mutants for membrane barrier defects using vancomycin. *A*. CFT073 and EC958 WT and FAS II mutants are cultured on LB-Lennox agar containing 50 *µ*g/mL vancomycin or DMSO carrier control. *B*. Complementation of defective membrane barrier in CFT073 and EC958 Δ*fabH* using plasmid-borne *fabH* expressed under arabinose control. Overnight cultures normalised to OD_600_ 1.0 were serially diluted to 1E-6, and 5 *µ*L of each dilution was spotted platted onto plates containing DMSO, 50 *µ*g/mL vancomycin or 100 *µ*g/mL fusidic acid. Where appropriate, 1% D-glucose or 50 mM L-arabinose were supplemented to suppress or induce the expression of plasmid-encoded *fabH*, respectively. The plate images shown are representative of at least three independent experiments.

As expected, WT strains of EC958 and CFT073 survived low-dose vancomycin, a Gram-positive antibiotic that is normally ineffective against Gram-negative bacteria as the OM prevents penetration to reach its peptidoglycan target. In contrast, CFUs of the Δ*fabH* mutants were diminished by 5-logs when exposed to 50 *µ*g/mL vancomycin (Fig. 1A). Likewise, we also observed similar increased sensitivity to fusidic acid, a hydrophobic antibiotic, that like vancomycin, is normally only effective against Gram-positive bacteria (Fig 2B). We then introduced the *fabH* gene into the Δ*fabH* mutants on the low copy-number and tightly-controllable pBAD322G vector under P*ara* control (28). Intrinsic resistance to both vancomycin and fusidic acid was fully restored to WT levels upon induction of FabH expression, but not when transcription was catabolically repressed (Fig. 1B). We also noticed a similar susceptibility pattern in laboratory K-12 strains; thus, the cryptic OM of Δ*fabH* strain is not restrictive to UPEC but generally applies to most, if not all, *E. coli* strains (data not shown).

**Fig 2.**
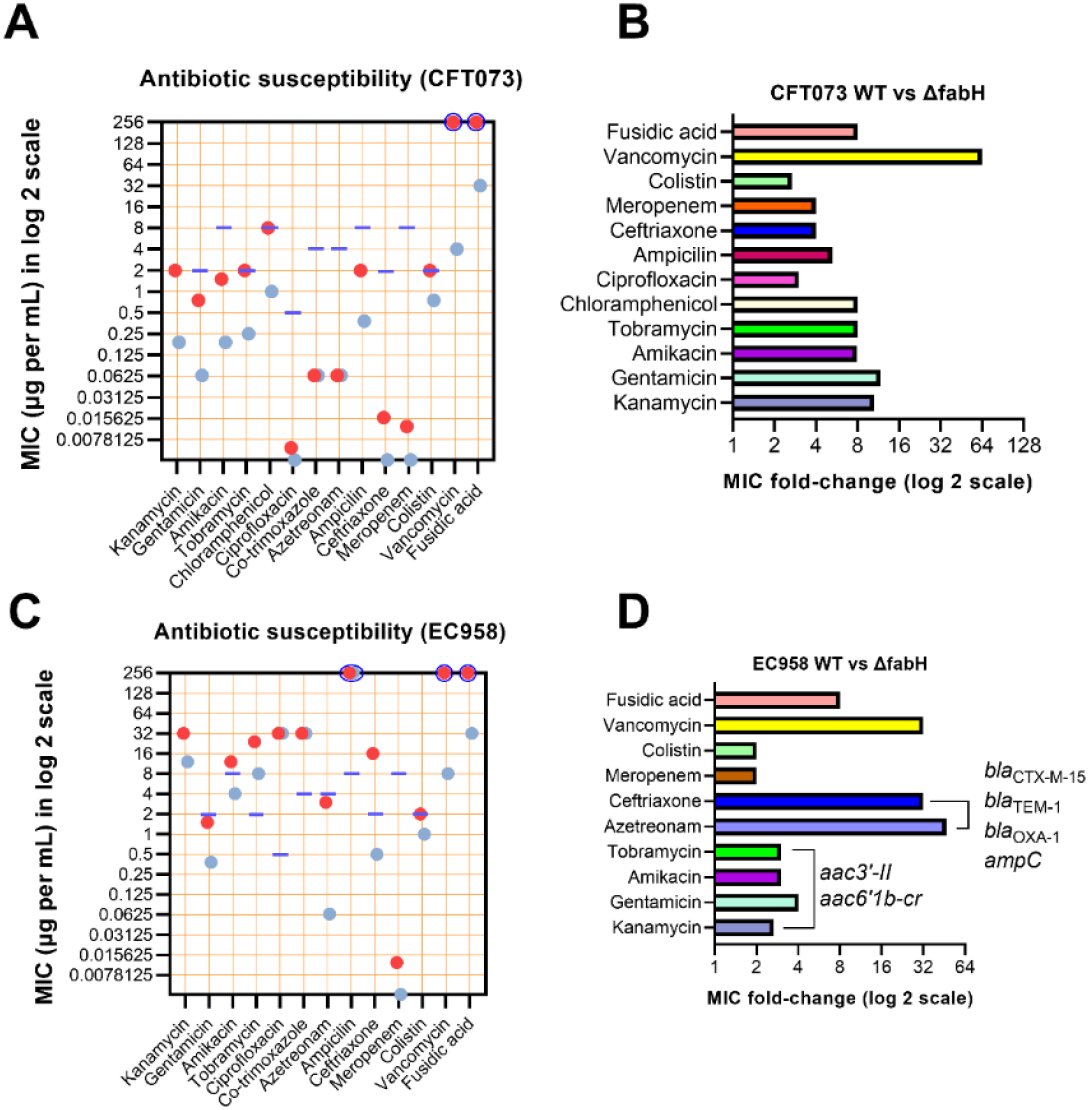
Loss of KAS III activity potentiates UPEC killing by a wide array of antibiotics. *A*, Individual antibiotic MICs for CFT073 WT (red dots) and Δ*fabH* strains (blue dots); *B*, Antibiotic MIC fold-changes (improvement in potency) for the CFT073 Δ*fabH* mutant relative to the WT. *C*, Individual antibiotic MICs for EC958 WT (red dots) and Δ*fabH* (blue dots); *D*, Antibiotic MIC fold-changes (improvement in potency) for the EC958 Δ*fabH* mutant relative to the WT. The purple lines in (A) and (C) mark current clinical resistance MIC cut-offs for each antibiotic as listed in the European Committee on Antimicrobial Susceptibility Testing Tables (30). MIC values that exceeded the testing range of the Liofilchem^®^ MIC strips are indicated by a blue circle that casts the red/blue dot(s).

### Loss of FabH promotes UPEC killing by several antibiotics

We hypothesised that the defective OM allows the efficient penetration of antibiotics and potentiates their inhibitory effect. To assess this, we determined the MIC of Δ*fabH* strains to a wide range of antibiotics (Fig 2). As expected, fusidic acid and vancomycin that cannot penetrate the Gram-negative cell envelope displayed an MIC of > 256 *µ*g mL^-1^ against CFT073 WT (Fig 2A). In marked contrast, for CFT073 Δ*fabH* the vancomycin MIC was reduced by >64-fold, whereas for fusidic acid, a >8-fold MIC reduction was observed (Fig 2A and 2B). Antibiotics representing the β-lactam, aminoglycoside, phenicol, polymyxin, and quinolone/fluoroquinolone classes were also tested. A >8-fold increase in susceptibility to both aminoglycosides (kanamycin, gentamicin, amikacin, and tobramycin) and phenicol (chloramphenicol) was observed. Moreover, the MIC values of ceftriaxone, meropenem, colistin, and ciprofloxacin against the isogenic CFT073 Δ*fabH* mutant were further decreased by 2- to 4-fold (Fig 2A and 2B), despite these antibiotics already being active at sub-*µ*g per mL concentrations against the susceptible CFT073 strain. Interestingly, we found CFT073 to be phenotypically resistant to tobramycin, chloramphenicol, and colistin, although it lacks recognisable antibiotic-resistant determinants (29).

The restored susceptibility to three antibiotics (tobramycin, chloramphenicol and colistin) in CFT073 Δ*fabH* prompted us to test if antibiotic efficacy could be restored against resistant strains. EC958 is an MDR clinical isolate of the globally disseminated ST131 lineage, carrying resistance genes to several antibiotic classes, including β-lactams (*bla*_CTX-M-15_, *bla*_TEM-1_, *bla*_OXA-1_ *bla*_CMY23_ and *ampC*), aminoglycosides (*aac3’-II* and *aac6’1b-cr*), and sulfonamides (*dhfrVII, aadA5, sul1*), in addition to two chromosomal mutations (S83L and D87N mutations in the *gyrA* gene) that confer fluoroquinolone resistance (23, 24, 31).

We attributed the very high co-trimoxazole and ciprofloxacin MIC values (> 32 *µ*g/mL) of EC958 WT and Δ*fabH* to the sulfonamide and quinolone resistance determinants present in the strain (Fig 2C). Like in CFT073, loss of *fabH* rendered EC958 susceptible to vancomycin (> 32-fold change in MIC), fusidic acid (> 8-fold), and colistin (2-fold) (Fig 2D). Moreover, relative to the WT, EC958 Δ*fabH* was >2-4-fold more susceptible to all four tested aminoglycosides (note that for EC958 WT, the kanamycin MIC exceeded the test range of the MIC strip, so the reported fold-change is likely an underestimate; Fig 2C and 2D). Importantly, amikacin susceptibility was restored in the EC958 Δ*fabH* mutant (Fig 2C and 2D), despite the presence of the aminoglycoside resistance genes (23, 31). We next compared the susceptibility of EC958 WT and Δ*fabH* strains to ampicillin, ceftriaxone, and aztreonam. As an extended-spectrum β-lactamase-positive (ESBL-positive) isolate, EC958 was found to be resistant to ceftriaxone (16 *µ*g/mL) and near intermediate-resistant to aztreonam (3 *µ*g/mL) (Fig 2C). For EC958 Δ*fabH*, we observed enhanced susceptibility relative to the WT. The MIC values for ceftriaxone and aztreonam were reduced by 32-fold, and ∼ 47-fold, respectively (Fig 2D). The extent of this MIC reduction rendered EC958 Δ*fabH* clinically susceptible to these otherwise ineffective antibiotics (with MICs well below the sensitivity values reported for Enterobacterales (30)).

The localisation and folding of outer membrane proteins can be impacted by compositional changes in the bacterial membrane envelope (see recent review by J. E. Horne et al. (32) for detailed information). Therefore, in addition to the severed permeability barrier, efflux pumps may be impacted in Δ*fabH*, and as such, partially contribute to the observed antibiotic hypersensitivity. We probed the efflux activity of WT and Δ*fabH* strains of both CFT073 and EC958, using intracellularly accumulated ethidium bromide (Fig 3). A Δ*tolC* mutant was constructed in EC958 as an efflux-defective control. As expected, ethidium bromide efflux was significantly delayed in this mutant (Fig 3). In contrast, ethidium bromide was rapidly expelled from both the WT control and the Δ*fabH* mutant (Fig 3). We conclude efflux activity is unaffected in UPEC Δ*fabH* strains.

**Fig 3.**
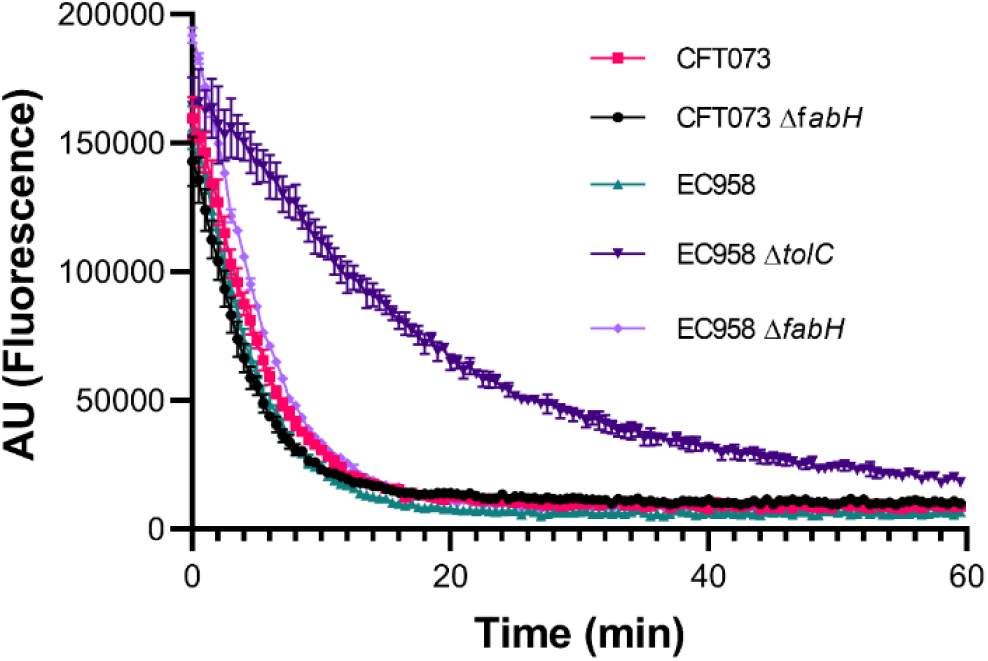
Efflux activity is unaffected in UPEC Δ*fabH* strains. Efflux kinetics of accumulated ethidium bromide were tracked in late phase bacterial cultures by excitation at 525 nm and emission at 615 nm. Experiments were performed in biological triplicates and average fluorescence readings (arbitrary units) taken every 30 secs are plotted with error bars showing standard deviation (SD) values.

### Loss of KAS III activity restores UPEC sensitivity to last line carbapenems

Intriguing, unlike other antibiotics of the β-lactam class, the Δ*fabH* strains were only twice more susceptible to meropenem than the WT strains (Fig 2D). Both UPEC strains lack genes for carbapenem resistance, so we introduced a *kpc2* containing plasmid (medium copy pSU2718 vector (33)) into the WT and Δ*fabH* strains to directly determine the impact of losing KAS III activity on carbapenem resistance.

Remarkably, the KPC-producing CFT073 Δ*fabH* strain displayed a reduced meropenem MIC by more than 21-fold relative to the WT strain (Fig 4). Similarly pronounced results were observed for three other carbapenem antibiotics, with the MIC of imipenem, doripenem and aztreonam reduced by 32-fold, > 31-fold, and > 42-fold, respectively (Fig 4B). All tested carbapenems (other than aztreonam) regained clinical potency against the KPC-producing UPEC Δ*fabH* strain. Overall, the loss of KAS III activity rendered the resistant parent strain remarkably susceptible to last-line carbapenems, with MIC values 4- to 10.5- fold lower than the current European Committee on Antimicrobial Susceptibility Testing resistance cut-off values for Enterobacterales (30).

**Fig 4.**
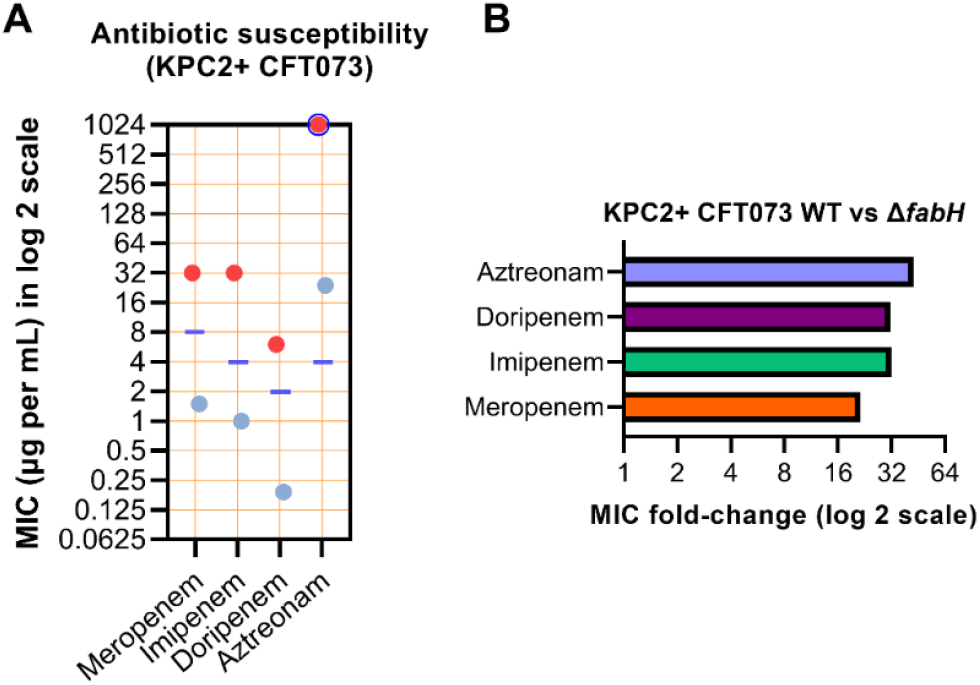
Carbapenemase-producing UPEC ΔfabH show restored susceptibility to carbapenem antibiotics. A. MIC values for four carbapenem antibiotics tested against KPC-producing CFT073 WT (red dots) and isogenic Δ*fabH* (blue dots). The purple line indicates the latest current resistant MIC cut-off value for each antibiotic listed in the European Committee on Antimicrobial Susceptibility Testing Tables (29). For MIC values that exceeded the testing range of the Liofilchem® MIC strips, a blue circle is used to cast the red/blue dot(s). *B*. Carbapenem susceptibility MIC fold-change (improvement in potency) for the KPC2+ CFT073 Δ*fabH* mutant relative to WT.

Prompted by the enhanced vulnerability of Δ*fabH* strains to last-line β-lactams and considering the severe OM defects observed in the UPEC strains lacking KAS III activity, we hypothesised that periplasmic β-lactamases may escape from the cell through the compromised OM, thereby reducing the periplasmic concentration of these protective enzymes in the previously resistant strain. To test this tenet, we collected cell-free media from the late log phase cultures of KPC2+ CFT073 WT and Δ*fabH* strains. Medium harvested from the KPC2+ Δ*fabH* culture, but not KPC2+ WT, rescued the growth of carbapenem-susceptible *E. coli* K-12 MG1655 on LB-Lennox agar containing 4 *µ*g/mL meropenem (Fig 5A). Proteinase K treatment of this growth medium reversed the growth rescue of MG1655. In contrast, heat treatment only marginally reduced the growth of MG1655 on meropenem LB-Lennox (Fig 5A). This observation is consistent with the previously reported thermostability of KPC2 (34).

**Fig 5.**
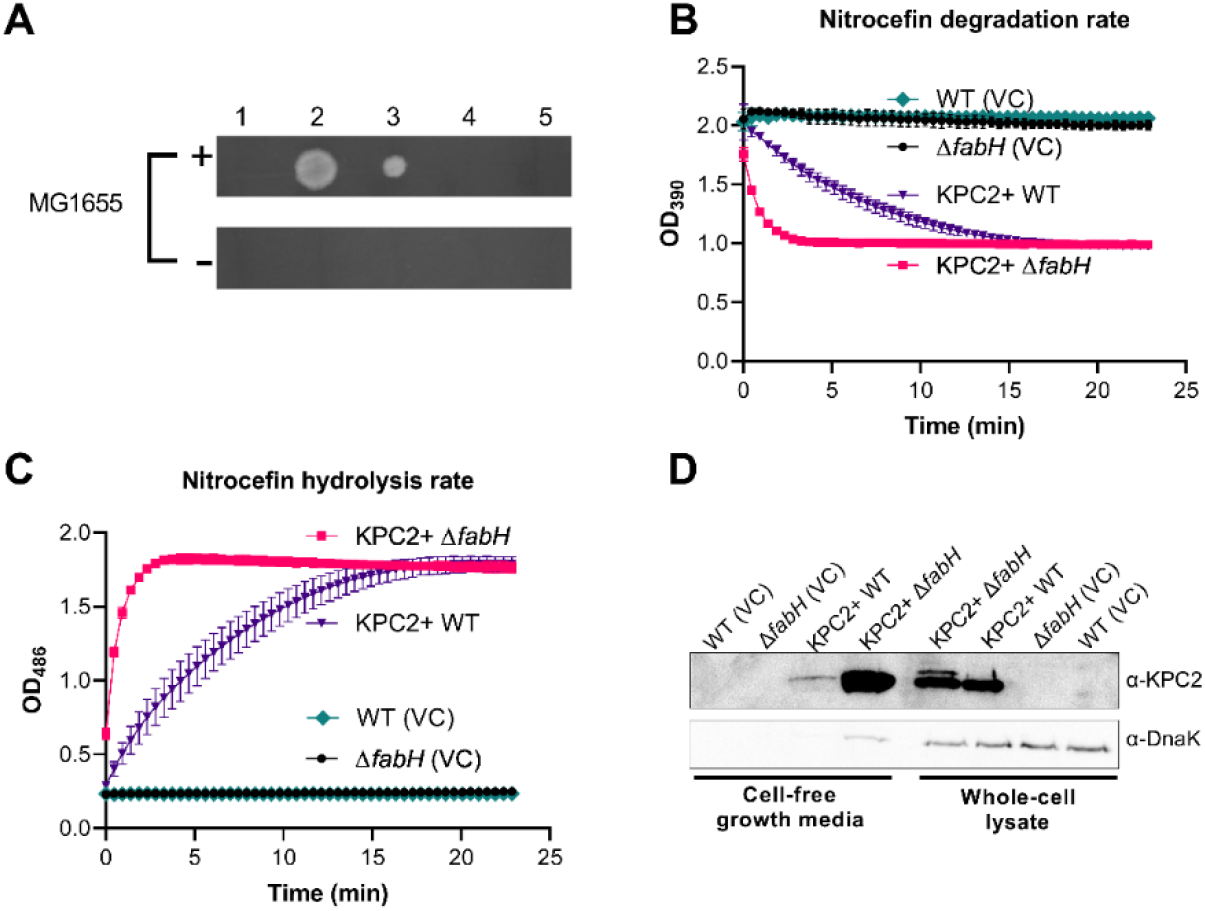
Evidence for leakage from the CFT073 Δ*fabH* severely compromised outer membrane. *A*, KPC2+ Δ*fabH* cell-free growth medium rescues MG1655 growth in meropenem containing agar. 1, Growth medium harvested from KPC2+ WT; 2, Growth medium harvested from KPC2+ Δ*fabH*; 3, Heat-treated KPC2+ Δ*fabH* growth medium; 4, Proteinase K treated KPC2+ Δ*fabH* growth medium; 5, LB-Lennox media control. Cell-free growth media were applied to meropenem plates swabbed with MG1655 (+) or no bacteria (-); *B*, Nitrocefin degradation assays using cell-free growth media recovered from KPC2+ CFT073 WT and Δ*fabH*; *C*, Nitrocefin hydrolysis assays using KPC2+ CFT073 WT and Δ*fabH* growth medium; *D*, Western blot of KPC2 present in cell-free growth media and whole-cell preparations; Data from three biological replicates are shown in panels B and C as means ± SD. Images shown in panels A and D are representative of three biological replicates.

We next diluted cell-free culture media to assay β-lactamase activity with nitrocefin, a chromogenic cephalosporin substrate (Fig 5B and 5C) (35). Growth media harvested from KPC2+ Δ*fabH* exhibited significantly stronger β-lactamase activity than that of the KPC2+ WT (Fig 5B and 5C). To confirm that KPC2 was present in significant amounts in the Δ*fabH* cell-free culture media, proteins from the supernatant were concentrated and analysed by Western blot using anti-KPC2 antisera. As expected, we detected KPC2 in whole-cell lysates of KPC2+ CFT073 but not in the vector control (Fig 5D). KPC2+ CFT073 Δ*fabH* had comparable amounts of KPC2 to the WT, although an additional higher molecular weight band consistent with premature KPC2 was also detected (Fig 5D).

In the cell-free culture supernatant fraction, we detected a weak signal for KPC2 in the KPC2+ CFT073 sample (Fig 5D). In striking contrast, very high amounts of KPC2 were observed in the KPC2+ Δ*fabH* culture medium sample (Fig 5D). A weak band corresponding to the cytoplasmic chaperone protein DnaK was also detected, indicating minor levels of cell lysis (Fig 5D). However, based on the very high KPC2:DnaK ratio in the sample compared to that of the whole cell (Fig 5D), we reasoned that the bulk of KPC2 detected in the cell supernatant likely escaped to the extracellular milieu through the severely compromised OM of Δ*fabH* intact cells.

### Loss of KAS III activity potentiates ceftriaxone treatment of ESBL+ UPEC biofilms and infected human bladder cells

UPEC form biofilms on biotic and abiotic surfaces that are recalcitrant to antibiotic treatment and aid bacterial survival inside bladder cells and on urinary catheters (36, 37). To investigate whether the enhanced antibiotic susceptibility of UPEC Δ*fabH* strains observed in planktonic drug sensitivity assays can be extended to improved activity against established biofilms, we grew EC958 WT and Δ*fabH* mature biofilms with both strains establishing comparable viable biofilm cell densities after 24 hours (Fig 6A). While biofilms formed by either strain had very high ceftriaxone resistance, a 4-fold reduction in the minimal biofilm eradication concentration (MBEC) was observed for Δ*fabH*, with a ceftriaxone MBEC of 512 *µ*g/mL for WT and 128 *µ*g/mL for Δ*fabH* (Fig 6B). Interestingly, residual EC958 WT cells remained viable even at the highest ceftriaxone dose tested (1024 *µ*g/mL), while no viable Δ*fabH* cells were detected in biofilms treated at and above 256 *µ*g/mL ceftriaxone, i.e., achieving complete biofilm eradication (Fig 6B).

**Fig 6.**
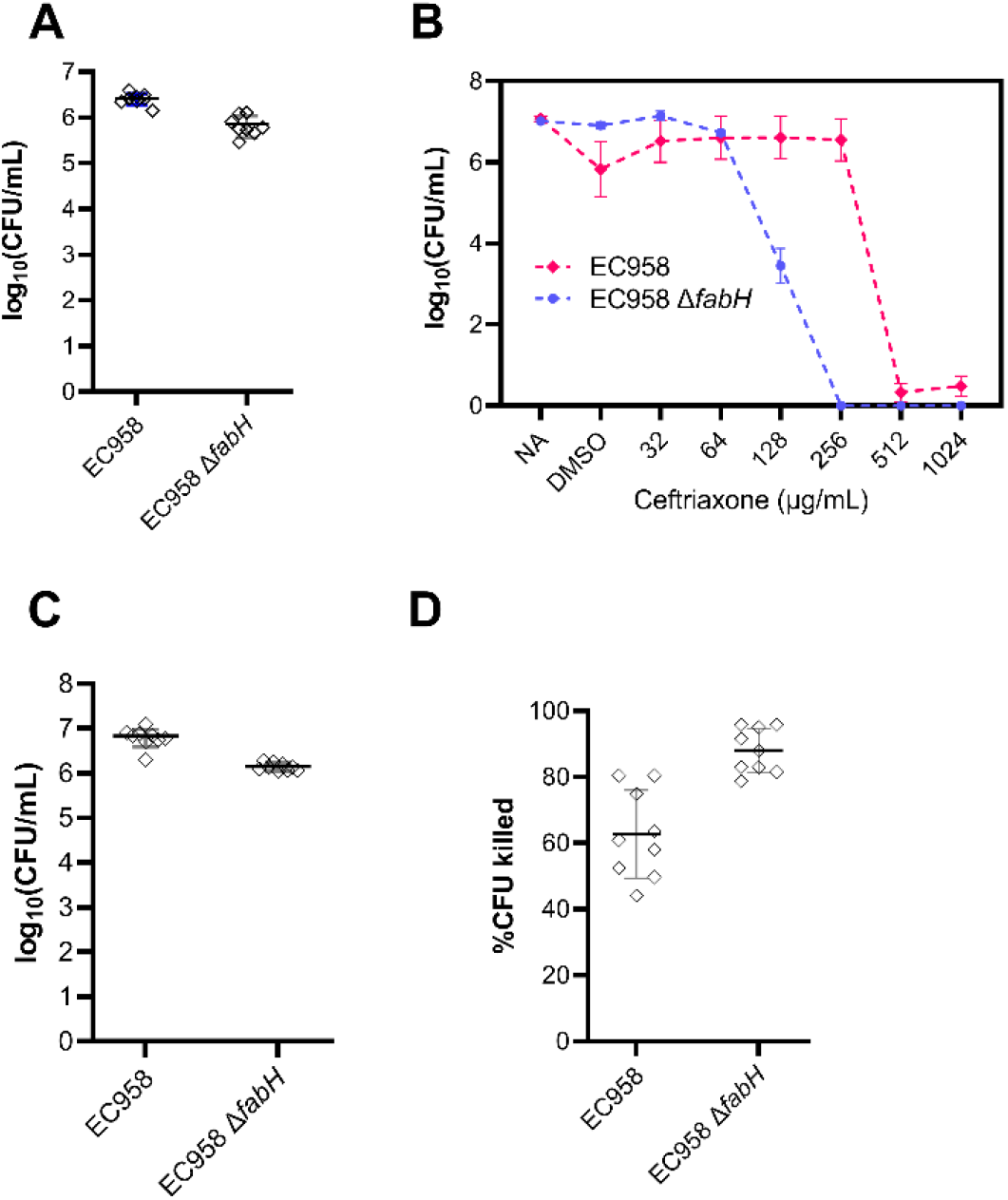
Preclinical evaluation of KAS III as a target aiding the eradication of UPEC biofilms and treatment of human bladder cell infection with antibiotics. *A*, EC958 WT and Δ*fabH* establish mature biofilms of comparable biofilm on the Calgary biofilm device; *B* ceftriaxone MBEC assessment of EC958 WT and Δ*fabH* biofilms, including untreated (NA) and drug carrier (DMSO) controls; *C*, Total adherent bacteria on T24 bladder cell monolayers infected for 24 hours at MOI 10 with EC958 WT and Δ*fabH*; *D*, Reduction in viable CFU recovered from UPEC infected T24 monolayers following one-hour treatment with ceftriaxone (8 *µ*g/mL).

We next tested if ceftriaxone is effective in treating established human cell infection by EC958 – a strain that is clinically resistant to this antibiotic. Infection of human T24 bladder cell monolayers with EC958 WT and Δ*fabH* at a multiplicity of infection (MOI) of 10, resulted in high level adhesion (>10^6^ CFU per monolayer) by both strains, albeit at slightly lower levels for Δ*fabH* (Fig 6C). Subsequent one-hour treatment of infected monolayers with 8 *µ*g/mL ceftriaxone, eliminated ∼ 90% of Δ*fabH* from the monolayer, while only a < 65% reduction was observed for WT (Fig 6D). Taken together, our preclinical data on improved antibiotic activity against resistant *E. coli* in planktonic, biofilm and cell infection models support the tenet that FabH constitutes a promising antimicrobial target for reviving failing last-resort antibiotics.

## DISCUSSION

AMR is one of the top ten global public health threats of the 21^st^ century, and tackling this ‘invisible pandemic’ constitute a current priority (38-40). A recent report highlights that 1.27 million deaths in 2019 were directly attributable to resistant bacterial infections, with *E. coli* the leading cause (6). Tactics to extend the life of existing antimicrobial agents or expand their spectrum are becoming a necessity in light of the dwindling antibiotic discovery pipeline, as new antibiotics are sparse. As an evolving permeability barrier, the OM is instrumental in developing antibiotic resistance in Gram-negative bacteria (9, 41). In recent years, there has been a resurgence of interest in exploring the disruption of the OM as an antimicrobial tactic (42-45). FAs are the common building blocks for structural lipids in the cytoplasmic membrane and the OM, thus their biosynthetic pathways present attractive antibiotic targets (46). Here we showed that the loss of KAS III incapacitates the OM to drastically improve antibiotic activity against clinical *E. coli*, even in the presence of acquired antibiotic resistance determinants.

Carbapenem is one of the last-resort antibiotics used to treat infections caused by drug-resistant Gram-negative bacterial infections. Unfortunately, resistance to carbapenems is increasingly prevalent in *E. coli* and other members within the Enterobacteriaceae family (47). We showed that targeting FabH can deactivate the antibiotic protection offered by advanced β-lactamases and carbapenemase to clinical *E. coli* strains. Another attractive prospect of targeting FabH (or other similar membrane perturbation tactics), is expanding the range of drugs with physicochemical properties amenable to Gram-negative entry, potentially expanding the spectrum of activity of many Gram-positive antibiotics.

FabH was previously thought to be indispensable in *E. coli* (48). However, this essentiality is bypassed by the product of the *yiiD* gene (renamed MadA) (49). The bypass mechanism involves the decarboxylation of malonyl ACP to produce acetyl ACP (50). Unlike acetyl CoA, which can only be integrated into the initiation step of FA biosynthesis by FabH, this substrate can be used by FabB/FabF to bypass the loss of KAS III activity in *E. coli* (50). However, the supply of FA to integrate into membrane biogenesis by the MadA bypass is highly restrictive, and as such, earlier *E. coli* K-12 studies reported that Δ*fabH* cells are tiny (26, 27). This observation had been instrumental in developing the lipid-centric view that FA availability sets the capacity of cell envelope to dictate cell size (27). Intriguingly, both CFT073 and EC958 Δ*fabH* cells have comparable size to their corresponding WT cells, though at the expense of cell division rate (Supplemental material, Fig S2). Together, these findings suggest that an additional layer of regulation might be employed in the two UPEC strains, allowing them to reach a destined cell envelope capacity despite severe FA starvation. Nonetheless, we note that antibiotic potentiation of Δ*fabH* is unrelated to cell size, given K-12 Δ*fabH* cells with reduced size also displayed increased sensitivity to hydrophobic antibiotics (26), and we also observed a 42-fold and 6-fold potentiation to vancomycin and fusidic acid in K-12 strains lacking KAS III (data not shown).

Two recent studies had linked FA starvation to increased antibiotic tolerance (51, 52). Sub-lethal inhibition of FA biosynthesis either by blocking the chain elongation steps catalysed by FabB/FabF or the enoyl ACP reductase FabI activates the (p)ppGpp synthetase, RelA, which overproduces ppGpp, the effector molecule of stringent bacterial response (51, 52). High ppGpp level inadvertently drives *E. coli* and several other species to reach an antibiotic tolerant state (51-57). The loss of KAS III also triggers the overproduction of ppGpp (26). In fact, the production of the stringent response alarmone is necessary for the survival of Δ*fabH*, as ppGpp is required to drive the transcription of MadA that partially substitutes FabH (49). Yet, in multiple *E. coli* lineages, Δ*fabH* shows hypersensitivity towards a broad panel of antibiotics.

Given the extent to which the cell envelope is disrupted in Δ*fabH*, allowing leakage of large periplasmic contents (Fig 5D), presumably, this damage alone is sufficient to overcome antibiotic tolerance induced by the stringent response. Nonetheless, to our surprise, the strain that possessed such a cryptic cell envelope remained highly stable, demonstrated by our failed previous attempts to isolate the then unresolved functional bypass of KAS III activity through permissive conditions (unpublished, Hong and Cronan). The mystery may lie in the multifaceted network regulating FAS II (58, 59) and other related pathways to compensate for defects in membrane biogenesis (60).

## MATERIALS AND METHODS

### Bacterial strains and growth

The strains and plasmids used in this study are described in Table 1. The *E. coli* strains were grown at 37 °C in Lysogeny broth (LB)-Lennox unless otherwise indicated. For solid media, 15 g/L bacteriological agar was added. Ampicillin was used at 100 *µ*g/mL, chloramphenicol at 17 *µ*g/mL, and gentamicin at 10 *µ*g/mL.

**Table 1.**
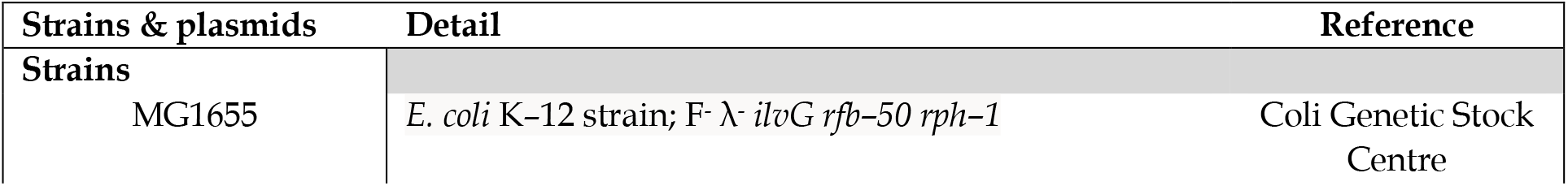
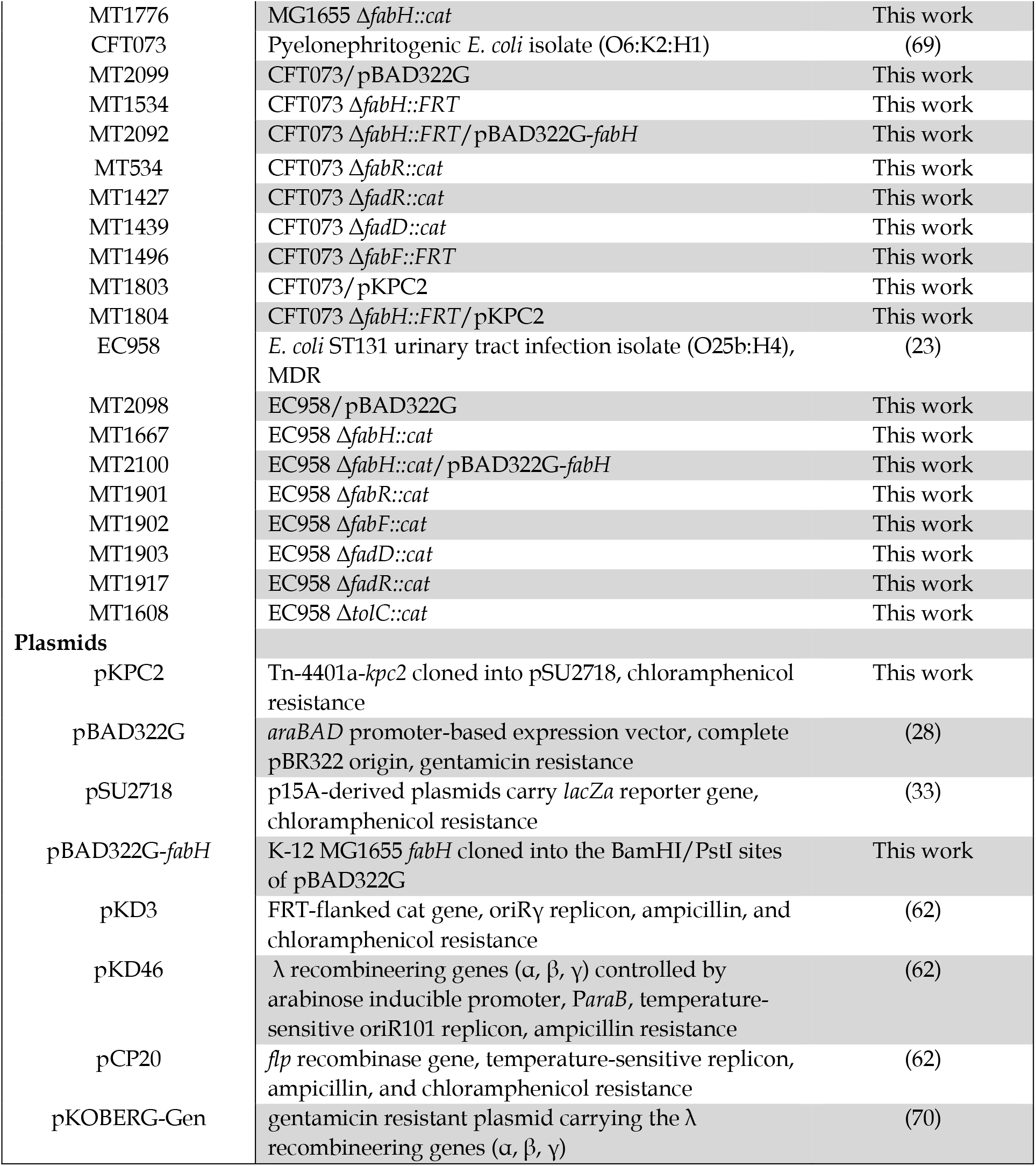
Strains and plasmids used in this work.

### Cloning and genetic manipulations

The *fabH* gene was amplified from the genomic DNA of the K-12 strain MG1655 and cloned into the BamHI and PstI sites in the pBAD322G vector (28). pKPC2 was constructed by amplifying Tn4401a-*kpc2* from the genomic DNA of a clinical *K. pneumoniae* strain JIE2709 (61), and cloned into PstI and KpnI sites in pSU2718 (33). Lambda recombination competency was achieved by adding temperature-sensitive plasmids, either pKD46 (62) or pKOBERG-Gen (63), to the manipulating strain. Transformation and lambda recombination were performed as previously described (64). Temperature-sensitive pCP20-Gent carrying FLP recombinase was used to remove the *cat* marker wherever applicable. For the construction of EC958 Δ*tolC::cat*, lamda-RED recovery culture post electrotransformation were platted onto low-dose chloramphenicol (6.5 *µ*g/mL chloramphenicol) LB-Lennox plate, and isolated and screened from thin bacterial lawn. Oligonucleotide sequences used for gene replacements and cloning were included in Supplemental material (Table S1).

### Outer membrane defect assay

Vancomycin can only transverses the Gram-negative OM when the permeability barrier is compromised (65). OM defect of the null mutants constructed was assayed using susceptibility of a dilution series from overnight grown cultures adjusted to OD_600_ 1.0 to subinhibitory concentrations of vancomycin similar to the method previously described (66, 67).

### Ethidium bromide efflux assay

Stationary cultures were diluted with fresh LB-Lennox containing 1 mg/L ethidium bromide and 20 *µ*g/mL carbonyl cyanide m-chlorophenylhydrazone. Cultures were incubated at 37 ºC until OD_600_ 0.7-0.8 with aeration. The cells pellets were harvested via centrifugation and washed with 4 × volumes of 1× PBS. The samples were then resuspended in 1× PBS and incubated for 30 min in a dark room at 5ºC. The samples were adjusted to OD_600_ 0.2 and dispensed in 150 *µ*L volumes in Greiner 96 well flat bottom plate; D-glucose (20% w/v stock) is then added at 0.1% concentration (12 *µ*L) to energise ethidium bromide efflux. Efflux activity was tracked over 60 min in CLARIOStar (BMG, Australia) at 37 ºC. The following instrumental settings were used, excitation at 525±15, emission at 615±20, auto 568.8 dichroic filter. The experiment was performed in biological triplicates.

### Determination of antibiotic minimal inhibitory concentrations

The specific bacterial strains were prepared and tested according to the MIC test strips (Liofilchem^®^) manufacturers’ instructions. Briefly, overnight cultures of the specific strains were diluted in 0.85 w/v % saline to give a final inoculum concentration of 1.5 × 10^8^ CFU mL^-1^. The inoculums were then used to inoculate Mueller-Hinton agar (Thermo Fisher, Australia) plates using a sterile swab. The MIC test strip was applied to the agar surface after the inoculum had dried. Plates were then incubated at 37 ºC for 20 hours. The MIC values were determined by observing where the relevant inhibition ellipse intersects with the MIC test strip.

### Determination of carbapenemase activity in cell-free growth media using MG1655 rescue

MT1803 and MT1804 overnight early stationary cultures were diluted 1:50 into fresh LB-Lennox containing chloramphenicol (17 *µ*g/mL) and grow to OD_600_ 0.8 at 37 ºC. Cell-free growth media were prepared by the filtration of culture supernatant of OD_600_ 0.8 cultures grown at 37ºC with aeration through a 0.22 *µ*m filter (3MM). The samples (5 *µ*L) were spotted onto LB-Lennox agar containing meropenem (4 *µ*g/mL) and had been swabbed with overnight cultures of MG1655 on the plate surface. Cell-free growth media treated with proteinase K at 37 ºC for 30 min, and heat (60 ºC for 30 min) treated cell-free growth media were also added to MG1655-swabbed LB-Lennox containing meropenem (4 *µ*g/mL) to serve as controls. LB-Lennox containing meropenem (4 *µ*g/mL) without MG1655 swab was also included as a control for the cell-free growth media. Plates were incubated at 37 ºC for 20 hours, and images were taken with BioRad GelDocXR+ system using the Image Lab software (v5.1; BioRad). The image shown is representative of three biological replicates.

### Assaying carbapenemase activity from cell-free growth media

MT1803 and MT1804 overnight early stationary cultures were diluted 1:50 into fresh LB-Lennox containing chloramphenicol (17 *µ*g/mL) and grow to OD_600_ 0.8 at 37 ºC. The cell-free growth media was prepared by filtering culture supernatant through a 0.22 *µ*m filter (3MM). The cell-free growth media were then diluted 1:1000 (180 *µ*L) and adjusted to 150 *µ*M nitrocefin to give a testing volume of 200 *µ*L. Nitrocefin degradation assays using cell-free growth media recovered from KPC2+ CFT073 WT and Δ*fabH* were then performed using KPC2+ CFT073 WT and Δ*fabH* growth medium, with two variables, degradation of nitrocefin and generation of products tracked at 390 nm and 486 nm on CLARIOStar (BMG, Australia) respectively, over time. Nitrocefin working stock was prepared at 1.5 mM in 50 mM phosphate buffer.

### Precipitation of extracellular proteins and Western blot

Five ml of Cell-free growth media were prepared by the filtration of culture supernatant of OD_600_ 0.8 cultures grown at 37 ºC with aeration through a 0.22 *µ*m filter (3MM). The sample is then precipitated with 1 ml of ice-cold 100% (w/v) trichloroacetic acid and incubated on ice for 10 min. The precipitated protein was harvested via centrifugation (20,000 ×*g*, 4 ºC, 10 min), washed once with 500 *µ*L ice-cold acetone, and dissolved in 100 *µ*L of SDS-PAGE sample buffer, with pH adjusted with 1M Tris-HCl, pH 9. Precipitated extracellular protein samples (10 *µ*L) and whole-cell lysates (10 *µ*L, derived from a 10× concentrate of OD_600_ 1.0 culture harvested simultaneously as cell-free growth media) were separated on 12% SDS PAGE. The samples were transferred to nitrocellulose membranes. Immunoblot analyses were performed with rabbit α-KPC2 (1:8,000) and mouse α-DnaK (1:10,000, Enzo Life Sciences, kindly gifted by Associate Professor Renato Morona), followed by goat α-rabbit Amersham ECL HRP-Conjugated Antibodies (1:20,000; Cytiva) for chemiluminescent detection using Pierce™ ECL Western blot substrate. Rabbit polyclonal α-KPC2 sera were raised by inoculation with three previously described KPC2 peptide epitopes [Epitope A: CFAKLEQDFGGSIGVYA; Epitope B: CLNSAIPGDARDTSSPRAVT; Epitope C: CVIAAAARLALEGLGVN) (68)] at the Walter and Eliza Hall Institute of Medical Research (peptides were synthesised by Mimotopes Pty Ltd). The developed blots were imaged in the BioRad GelDocXR+ system using the Image Lab software (v5.1; BioRad).

### Biofilm growth

Minimum biofilm eradication concentration (MBEC) and biofilm growth assays were performed in a Calgary biofilm device (CBD) (MBEC assay; Innovotech Inc., Canada). Overnight bacterial cultures in LB were diluted to 10^6^ CFU/ml and used to inoculate the plate with 130 μL of culture per well. The CBD was incubated for 24 hours with shaking (150 rpm) at 37 °C in 95% relative humidity. Following 24 hours of growth, biofilms were washed once in PBS to remove non-adherent cells and then sonicated for 20 min at 20 °C. At least three biological and two technical repeats per strain per experiment were serially diluted and spotted onto LB agar plates to determine viable CFU recovered from each peg biofilm. Plates were incubated overnight at 37 °C with colonies counted the following day to obtain log_10_(CFU/mL) values for each strain.

### MBEC assays

Following 24 hours of biofilm growth in the CBD, biofilms were washed once in PBS to remove non-adherent cells and the biofilm peg lid was transferred to a treatment plate containing varying concentrations of ceftriaxone disodium salt heptahydrate or dimethyl sulfoxide (DMSO) in Muller Hinton broth. Peg biofilms were incubated in the treatment plate for 24 hours and then sonicated and plated as per the biofilm growth assay. MBEC values were defined as concentrations with over 3 log_10_ reduction in CFU/mL.

### Epithelial cell infection assay and ceftriaxone treatment

Intestinal epithelial cells T24 (ATCC HTB4; DMEM) were maintained in McCoy media (Invitrogen) supplemented with 5% heat-inactivated foetal calf serum (Invitrogen). Bacterial strains were cultured under type 1 fimbriae enrichment conditions, as previously described (23). Infection assays were performed as previously described (63). Briefly, confluent cell monolayers were infected with strains at a multiplicity of infection (MOI) of 10 and incubated at 37 °C, 5% CO_2,_ for 1 hour. PBS washes (3×) were used to remove non-adherent bacteria. Ceftriaxone salt heptahydrate (in DMSO) was added to McCoy media and applied to the infected monolayer without disturbance. Antibiotic treatment was performed by incubation at 37 °C, 5% CO_2,_ for 2 hours. 1× PBS washes (3×) was used to remove non-adherent bacteria and antibiotic residue. Monolayers were lysed with 0.1% (v/v) Triton-X100, and lysates were serially diluted and plated onto LB agar to enumerate total adherent bacteria.

## ACKNOWLEDGEMENT

Rabbit α-EcDnaK is a generous gift from Associate Professor Renato Morona (University of Adelaide, Australia). This work was supported in part by Australian National Health and Medical Research Council Project Grant (GNT1144046), a Clive and Vera Ramaciotti Health Investment Grant (2017HIG0119), Australian Research Council (DE130101169) and a Georgina Sweet Award for Women in Quantitative Biomedical Science to MT and an Early Career Research Grant from the Institute of Health and Biomedical Innovations at the Queensland University of Technology, Australia, to YH. CLARIOStar high performance microplate reader (BMG, Australia) was sponsored by the Ian Potter Foundation. SH is the recipient of an Australian Government Research Training Program (RTP) Scholarship. BZ was supported by a China Scholarship Council (CSC) scholarship. MT received support from Queensland University of Technology through a Vice-Chancellor’s Research Fellowship.

## AUTHOR CONTRIBUTION

YH, JEC and MT contributed to project conception; YH, JQ and MT contributed to experimental design. YH, JQ, ADV, SH, and BZ conducted experiments, data collection, analysis, and interpretation; YH and MT supervised the study and obtained the funding. YH drafted the initial manuscript. YH and MT substantially revised the manuscript. JEC and MATB contributed to data interpretation and revised the manuscript. All authors approved the final manuscript.

